# Scalable, high-throughput isolation of extracellular vesicles using electrokinetic-assisted mesh filtration

**DOI:** 10.1101/2025.03.27.645682

**Authors:** KangMin Lee, Minju Bae, YongWoo Kim, SoYoung Jeon, Sujin Kang, Wonjong Rhee, Sehyun Shin

**Affiliations:** School of Mechanical Engineering, Korea University, Seoul, Republic of Korea; Department of Micro-Nanosystem Technology, Korea University, Seoul, Korea; Engineering Research Center for Biofluid Biopsy, Seoul, Korea; Department of Bioengineering and Nano-Bioengineering, Incheon National University, Incheon, Republic of Korea

**Keywords:** EV, isolation, scalable, high-throughput, electrokinetic, ExoFilter

## Abstract

As extracellular vesicles (EVs) are increasingly recognized for their superior functions for therapeutics, the need for large-scale EV isolation technology is becoming more critical for clinical and industrial applications. Most existing EV isolation methods are optimized for small-scale laboratory samples, limiting their efficiency and scalability for large-scale production. Here, an electrokinetic-assisted filtration system (ExoFilter), which introduces charge interaction into physical mesh flow filtration, is proposed as a new candidate to address the challenges of scalable EV isolation. The hybrid filtration system demonstrates outstanding high-throughput EV isolation performance (a flux of ∼750 mL/min) using only a coarse physical filter, by electrokinetically arresting EVs flowing through the filter lattice. Furthermore, the recovery efficiency of ExoFilter, analyzed based on the ELISA results, was found to be approximately 98%, demonstrating the filter’s exceptional efficiency in EV isolation. Additionally, ExoFilter enables the rapid isolation of EVs from small samples as little as 200 µL, facilitating quick and easy blood-based EV research. Furthermore, low-molecular-weight albumin from plasma samples was effectively removed. The high-throughput and high-efficiency characteristics of ExoFilter make it well-suited for scalable EV production, offering greater convenience for various clinical applications.

**Highlights:**

- Electrokinetic-assisted mesh filtration (ExoFilter) enables scalable and rapid isolation of extracellular vesicles (EVs).
- A high throughput of ∼750 mL/min is demonstrated while maintaining high yield and purity.
- ExoFilter effectively removes albumin contaminants from EVs through size-exclusive electrokinetic-assisted mesh filtration.
- Efficient EV isolation performance is achieved for human plasma, saliva, urine, and cell culture media.

**Graphical abstract:** 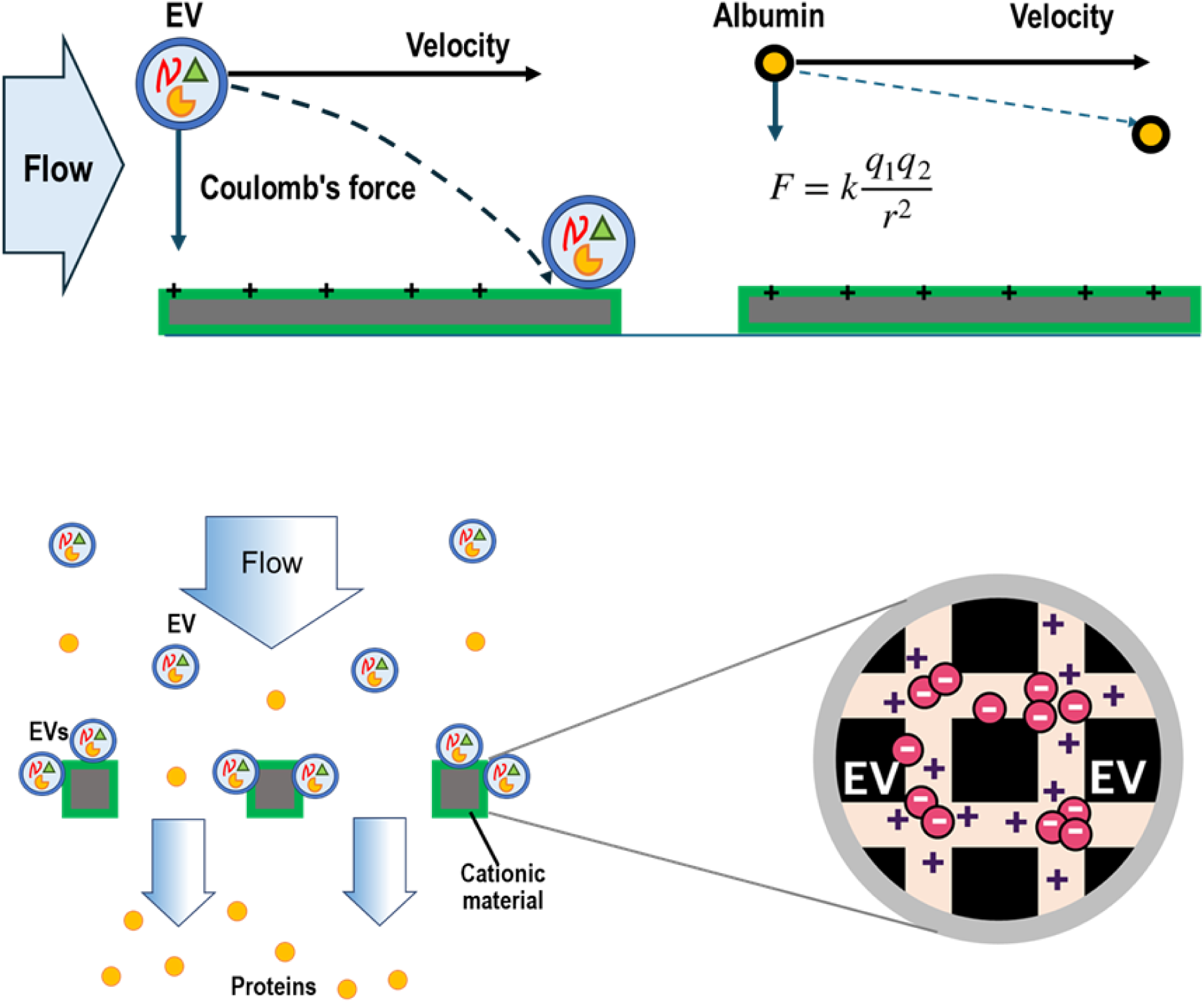

## 1. INTRODUCTION

Extracellular vesicles (EVs) have emerged as excellent mediators of cell-to-cell communication due to their ability to cross biological barriers [(Pitt et al., 2016), (Nawaz et al., 2018), (Wiklander et al., 2019)]. These vesicles, secreted by various cells, carry a rich assortment of biomolecules, including proteins, lipids, RNA, DNA, and various nucleic acids and proteins. These cargoes can be specifically transferred to target cells via interactions between EV surface molecules and target cells, ensuring precise targeted delivery. Due to their natural origin and ability to deliver bioactive molecules, EVs are being explored as therapeutic delivery vehicles [(Akers et al., 2013), (Vader et al., 2016), (Momen-Heravi et al., 2018)]. EVs are now highly utilized to carry specific therapeutic agents, such as drugs or genetic material, making them promising tools for targeted therapy. Recently, engineered EVs have opened exciting advancements in therapeutic delivery and molecular communication, allowing vesicles to carry specific therapeutic cargoes, such as proteins, nucleic acids, or drugs, thereby enhancing their potential for targeted therapy.

As the application of extracellular vesicles (EVs) advances toward practical and industrial use [(Raposo & Stoorvogel, 2013), (Cardoso et al., 2021)], there is a growing demand for innovations in large-scale EV isolation techniques. High-throughput and efficient isolation methods are crucial for producing the quantities of EVs required for clinical and industrial purposes (Paganini et al., 2019). However, current isolation techniques, which were primarily designed for small-scale laboratory use, face significant challenges when scaled up to larger volumes [(Gimona et al., 2017), (Stahl & Raposo, 2018), (Théry et al., 2018)]. Industry typically demands EV production at scales ranging from several liters to hundreds of liters per batch, which existing methods struggle to achieve efficiently. Emerging technologies such as microfluidics, continuous-flow ultracentrifugation, and tangential flow filtration (TFF) are being explored to overcome these obstacles, aiming to enable the consistent and efficient production of high-quality EVs (Stam et al., 2021).

The current state of technological advancement remains inadequate, highlighting the urgent need for further innovation to meet the growing demands of the industry and facilitate the successful commercialization of EV-based therapies and products. After reviewing previous research, the primary challenges in large-scale EV isolation include the difficulty in scaling up traditional methods while maintaining high yield, purity, and EV integrity, as well as the need for efficient processing within a reasonable time and cost (Kapoor et al., 2024). Additionally, achieving consistency and reproducibility across batches is crucial, particularly for clinical and industrial applications. Thus, an innovative approach is necessary to address these difficulties in large-scale EV isolation.

This study presents a new ultrafast scalable isolation technique, charge-based electrokinetic mesh filtration (ExoFilter), for EV isolation that overcomes the limitations of other methods. To validate the effectiveness of ExoFilter, we analyzed isolated EV samples according to established guidelines in the EV field, such as MISEV2023 (Welsh et al., 2024). The EV fractions were thoroughly characterized with a focus on the enrichment of EV-specific markers and the absence of non-EV proteins. The size distribution and concentration of EVs were analyzed using nanoparticle tracking analysis (NTA), while EV purity was assessed through standard protein detection methods, and their morphology was visualized using SEM and TEM. We compared the performance of ExoFilter with other methods, including UC, TFF, ExoQuick, and ExoPAS, confirming that ExoFilter provided superior yield and purity compared to existing technologies, while offering significant improvements in processing speed. Additionally, as proof of concept for large-volume sample extraction, we demonstrated results across various samples for 500 mL. These findings highlight that ExoFilter enables faster processing of large-volume samples, achieving better cost efficiency and a reduced environmental impact.

## 2 RESULTS

### 2.1 Operating principles of ExoFilter

This study developed a novel charge-based filtration method to isolate EVs for large sample volumes. As shown in Figure 1(a), ExoFilter employs a multi-layered mesh coated with cationic materials to selectively capture negatively charged EVs through electrokinetic interactions. The effectiveness of charge-based EV isolation has been demonstrated in our previous research [(Kim et al., 2020), (Kim & Shin, 2021)] as well as in other studies [(Deregibus et al., 2016), (Petga et al., 2024)]. As shown in Figure 1(b), EVs isolated with ExoFilter have a zeta potential of -15.5 mV, while the cationic nylon mesh shows +15.4 mV, resulting in strong electrostatic interactions that effectively bind EVs to the mesh surface. For comparison, plasma proteins such as albumin (−12.1 mV), γ-globulin (7.9 mV), and fibrinogen (−4.5 mV) are also presented. Negatively charged proteins remain difficult contaminants to remove using anion exchange methods like ExoFilter.

**Figure 1.**
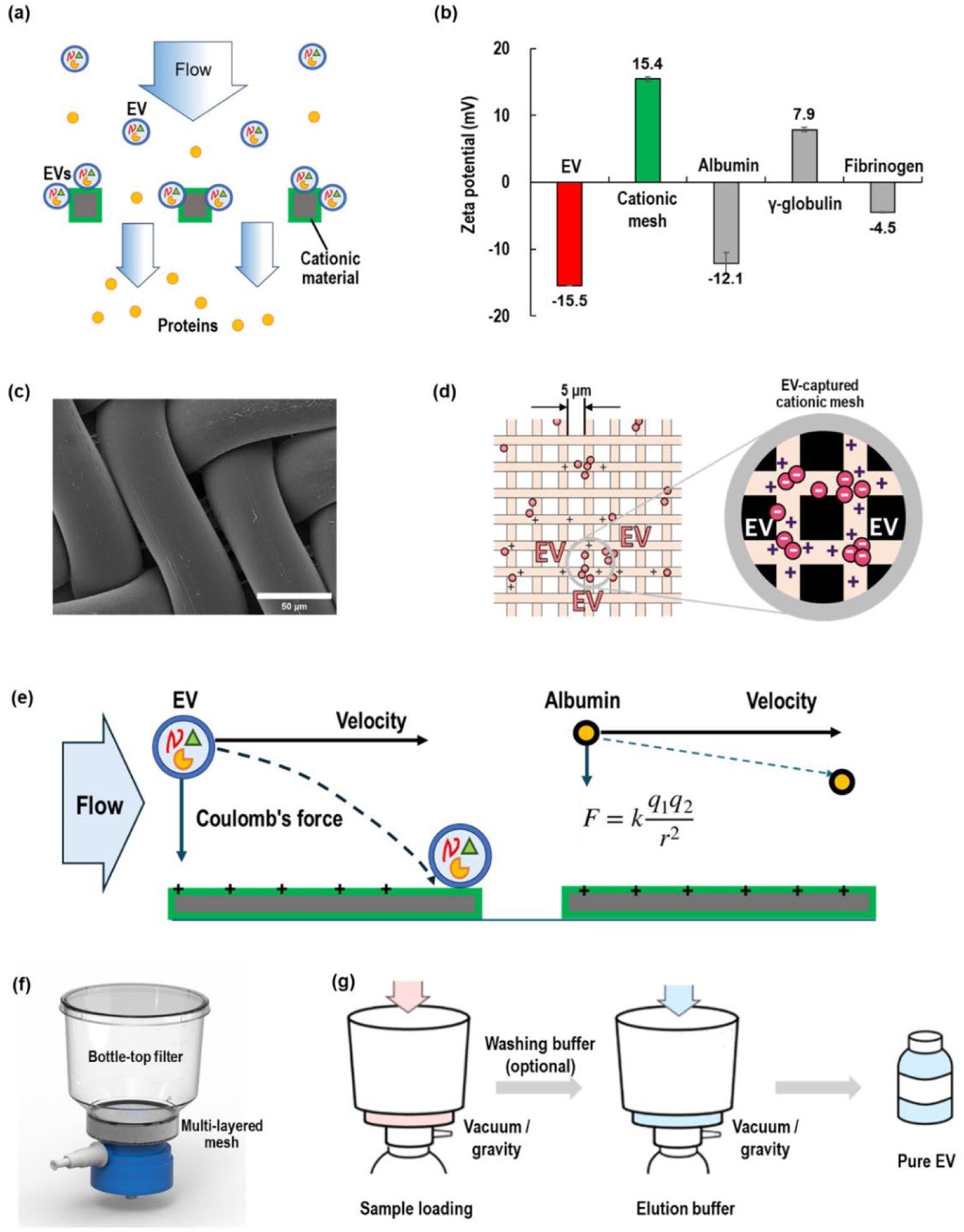
Schematics of EV isolation using electrokinetic-assisted mesh filtration (ExoFilter). (a) Mechanism of ExoFilter, (b) Zeta potentials of EV, cationic nylon mesh and plasma proteins (albumin, γ-globulin, fibrinogen), (c) SEM image of cationic mesh, (d) EVs-captured mesh via electrostatic interaction (mesh pore size, 1 um). (e) Electrokinetic-assisted filtration for EV and albumin, (f) Modified bottle-top filter with multi-layered cationic meshes, (g) Protocol of ExoFilter.

Figure 1(c) shows the Scanning Electron Microscope (SEM) image of the nylon mesh used in the ExoFilter. These mesh structures, featuring triangular channels with a base length of 20 µm and a height of 5 µm, allow fluids to pass smoothly while effectively capturing nano-sized particles such as EVs. The larger mesh pore size highlights that the present method is not reliant on size exclusion, addressing the scalability imitations of previous approaches. Owing to the and multi-stacked mesh and the structures of mesh pores, ExoFIlter provides a high surface area for maximum interaction with target particles, maximizing the efficiency of the ExoFilter. Figure 1(d) illustrates a schematic of the ExoFilter mechanism, demonstrating how negatively charged EVs are attracted and captured by the positively charged surface.

ExoFilter introduces charge interaction mechanism into physical mesh flow filtration, which is so called electrokinetic-assisted mesh filtration system. It is worth noting that the mesh filter is quite coarse and thus EVs can freely pass through filter pores if there is no charge-interaction. During the operation of ExoFilter, particles in a sample dominated by two physical force interactions between electrostatic forces and the inertia of flowing particles, as shown in Figure 1(e). One thing to note is that even for nanoparticles with the same zeta potential, as the size of the particle increases, the surface area also increases, resulting in a larger charge and consequently a stronger Coulomb force. Thus, when a highly charged large particle is located at a suitable distance from a surface with the opposite charge, the strong Coulomb force exerted on the particle causes it to follow a steep trajectory across the fluid streamlines toward the surface. In contrast, a smaller particle with a lower charge tends to deviate only slightly from its original streamline and cannot be captured on the mesh surface. This process, where larger charged particles are selectively separated while smaller ones remain in the fluid, can be described as a kind of size-exclusion electrokinetic-filtration (Lee et al., 2023).

Figure 1(f) depicts an example of the ExoFilter, modified from a standard bottle-top filter, where the original filter element is replaced with a multi-layered mesh. The original capacity of the bottle-top filter to accommodate 250 mL is retained, enabling the ExoFilter to efficiently process samples up to 1 L. The workflow is remarkably simple and time-efficient, consisting of just two straightforward steps: sample loading (with optional washing) and elution. First, a sample containing EVs is introduced into the ExoFilter. Using vacuum, the sample is quickly drawn through the filter. If necessary, a washing buffer can be applied to remove non-specifically bound proteins and contaminants. The washing buffer was prepared by adding acetic acid to water and adjusting the pH to 6. Finally, an elution buffer (NaCl 1 M) is used to release the captured EVs from the mesh, resulting in a final eluate enriched with purified EVs. Typically, the volume of the elution buffer used is approximately one-fifth of the sample volume, although it can be reduced to as little as one-tenth, depending on the requirements.

### 2.2 Characterization of EVs isolated with ExoFilter

Figure 2 provides a comprehensive validation of EVs isolated from plasma sample 1 mL using the ExoFilter, confirming their identity through morphology, size distribution, and characteristic EV markers. First, we present SEM and TEM images of EVs isolated from a blood sample using the ExoFilter. These EM images confirm the spherical morphology and phospholipid bilayer of the EVs. The NTA results show that the size distribution of EVs isolated by ExoFilter ranges from 30 nm to 300 nm, with a mode at 122 nm and a mean size of 141 nm.

**Figure 2.**
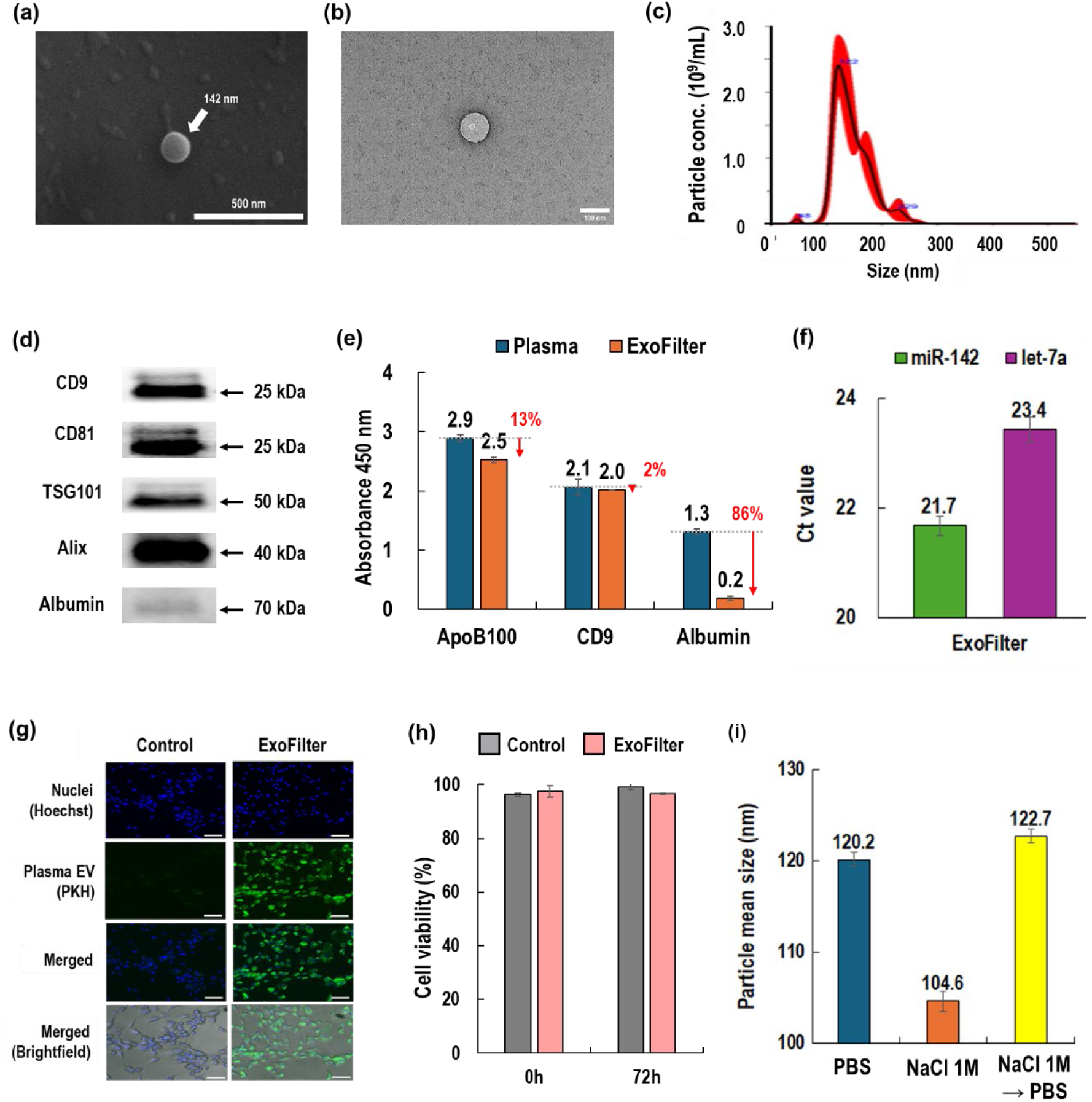
Characterization of EVs isolated from 1 mL of blood plasma using ExoFilter. (a) SEM image of EV, (b) TEM image of an eluted EV, (c) Particle size distribution and concentration of EVs (elution sample diluted 1/100) (d) Western blot assay for isolated EVs, (e) ELISA for isolated EVs, (f) Ct values of RT-PCR for miRNAs of isolated EVs, (g) Cellular uptake of EVs into human dermal fibroblasts (HDFs), (h) Cytotoxicity test of EVs in HDF after 72 hours, (i) Variation of particle size with elution buffers.

Through Western blot analysis, we detected the tetraspanin markers CD9 and CD81, along with internal proteins TSG101 and Alix (full Western blot image available in Figure S1). Additionally, albumin, the most abundant negatively charged plasma protein, showed a faint band of similar intensity to TFF (Figure S1), indicating successful depletion of low molecular weight proteins by ExoFilter. Furthermore, an ELISA assay confirmed the expression of surface markers CD9, which are distinct EV biomarker, while also detecting albumin and Apo B-100. Albumin, plasma protein contamination marker, was significantly reduced by approximately 86% after filtration with ExoFilter, as shown in Figure 2e. However, the presence of Apo B-100, a biomarker for VLDL and LDL, highlights a limitation of the electrokinetic-assisted mesh flow filtration, as it was not completely removed due to its size overlap with EVs.

Furthermore, we confirmed the presence of EV-derived miRNA through RT-PCR. After lysing the isolated EVs, RNA was extracted and amplified for two housekeeping genes (miR-142 and let-7a). The results demonstrated sufficient detection of these miRNAs, confirming that the particles isolated by ExoFilter are indeed plasma-derived EVs. Through cellular uptake experiments, the isolated EVs were effectively internalized by human dermal fibroblasts (HDFs), demonstrating successful uptake. Additionally, a 72-hour cytotoxicity test indicated that the isolated EVs did not induce significant cytotoxic effects, maintaining high cell viability and full functional bioactivity.

There was a potential osmotic pressure issue associated with using 1 M NaCl as an elution buffer. As shown in Fig. 2(i), EVs recovered with 1 M NaCl exhibited a reduction in particle mean size to 104.6 nm, compared to 120.2 nm when recovered with PBS. However, the particle concentration remained statistically unchanged (Figure S2). Notably, when EVs recovered with 1 M NaCl were exchanged with PBS, the particle mean size fully recovered to 122.7 nm. After elution with 1 M NaCl, the samples were stored at -80°C for a minimum of 1 week and up to 4 weeks prior to SEM and TEM image analysis. Additionally, cellular uptake and toxicity experiments were conducted. These findings suggest that the high concentration of 1 M NaCl may reduce particle size due to osmotic pressure. However, the structural integrity and functionality of EVs remain uncompromised, as the size reduction is reversible upon dilution with PBS.

### 2.3 ExoFilter optimization

To enhance the yield and purity of EV isolation using the ExoFilter, several key parameters, including the number of mesh layers and the effectiveness of the washing steps, were systematically optimized. Initial experiments for optimizing the number of mesh layers were conducted using a smaller ExoFilter designed for a 1 mL plasma sample in a spin-column format. Subsequently, to evaluate the effectiveness of the washing steps, an ExoFilter bottle-top filter was used to process the diluted plasma sample. Specifically, a 5 mL plasma sample was diluted 1:100 in PBS, yielding a total volume of 500 mL. This diluted sample was then eluted with 100 mL of 1 M NaCl. For the elution step, a volume equivalent to one-fifth of the sample volume was applied as the elution buffer.

#### Optimization of Mesh Layers

Although the final goal was to use a scalable sample volume, a 1 mL ExoFilter was fabricated and utilized for the optimization phase. Since the ExoFilter captures EVs via electrokinetics through meshes, an appropriate surface area of the mesh is essential for a given sample volume. Therefore, extraction efficiency is highly dependent on the surface area, which increases with the number of mesh layers. As shown in Figures 3(a) and 3(b), the spin column-type ExoFilter can be configured with varying numbers of mesh layers. In this study, we compared the efficiency of ExoFilters with 1, 10, 20, and 30 mesh layers.

**Figure 3.**
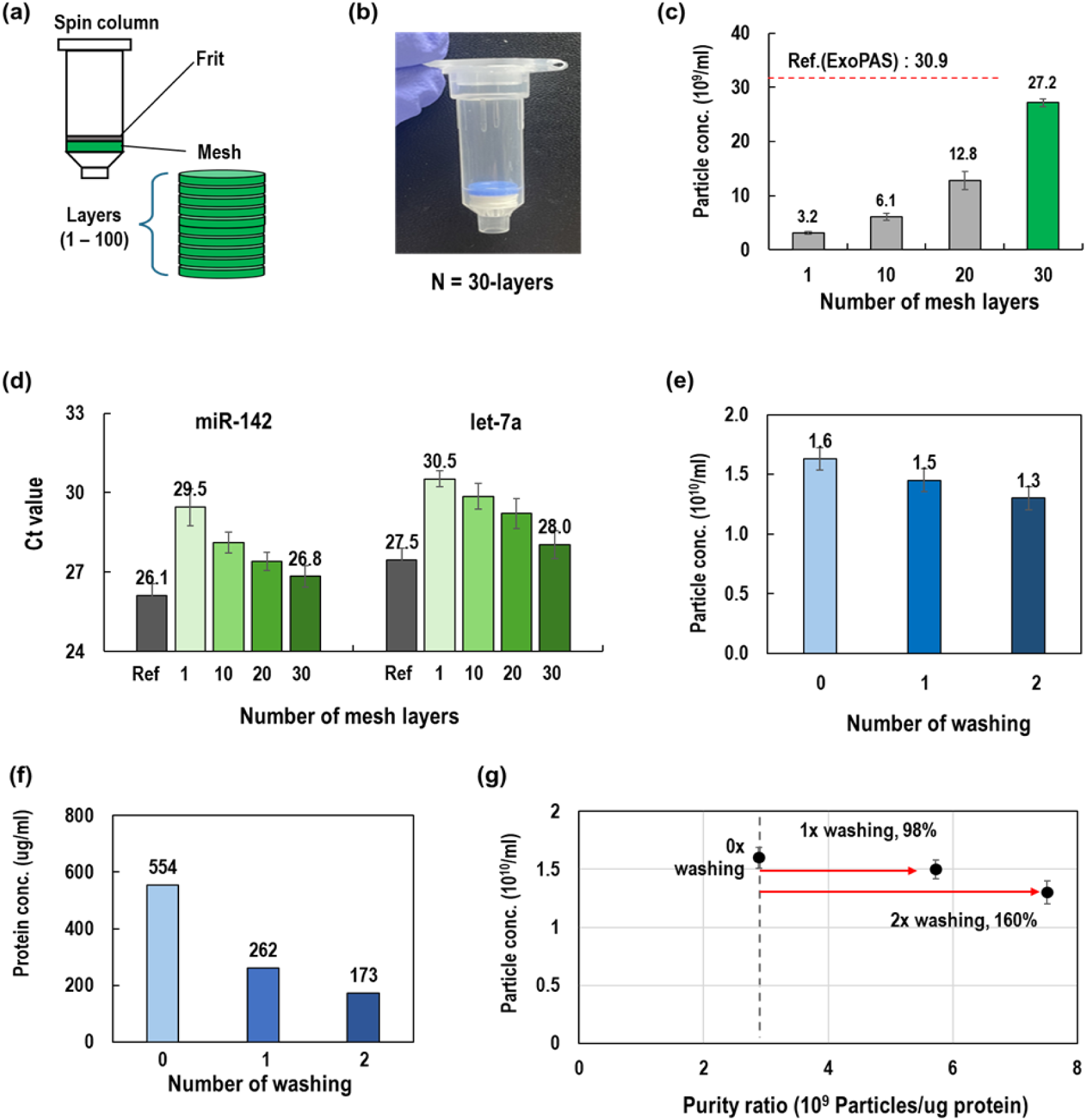
Optimization of the ExoFilter system using plasma (5 mL) diluted to a final volume of 500 mL (1/100 ratio). (a) Configurations of an ExoFilter with varying different number of meshes, (b) An photographic image of ExoFIlter with 30-layered meshes, (c-d) Effects of number of mesh layers on particle concentration (NTA) and microRNAs (RT-qPCR), (e-g) Effect of washing on particle concentration, protein concentration and purity ratio.

As the number of mesh layers increased, the concentration of extracted particles also increased proportionally, with the 30-layer ExoFilter achieving the highest extraction yield (Figure 3(c)). The extraction performance of the ExoFilter was approximately 90% of that of the ExoPAS method, which is remarkable considering that precipitation-based methods likely include many contaminants. To further validate the efficacy of the ExoFilter, we analyzed miRNAs, including the housekeeping genes miR-142 and let-7a, through RT-qPCR (Figure 3(d)). The cycle threshold (Ct) values of these miRNAs decreased as the number of mesh layers increased, indicating improved miRNA detection. Specifically, the Ct value for miR-142 decreased from 29.5 to 26.8, and for let-7a, from 30.5 to 28.0 with 30 mesh layers. The highest values of ExoFilter are nearly identical to those of ExoPAS, confirming similar results observed in the NTA results in Figure 3(c).

#### Optional washing process

After optimizing the number of mesh layers, we proceeded with optimization experiments for the washing step using large-volume samples. The a 500 mL plasma sample diluted 1:100 was used. The washing buffer was prepared by mixing acetic acid with deionized water and adjusting the pH to 6.0. The washing step was performed using 500 mL of the washing buffer. NTA results (Figure 3(e)) showed a modest reduction in particle concentration after one and two washing steps, with decreases of 9.4% and 18.7%, respectively, compared to the unwashed condition. However, protein contamination was significantly reduced with each additional washing step (Figure 3(f)). Specifically, protein concentration dropped from 554 µg/mL to 262 µg/mL (a 47.3% reduction) after 1x washing and further decreased to 173 µg/mL (a 68.8% reduction) after 2x washing.

These results underscore the effectiveness of the washing steps in removing non-specifically bound proteins and other contaminants. Considering the results from both NTA and BCA assays, the optional washing process led to minimal reduction in particle recovery while significantly reducing protein contamination. This outcome is further supported by the purity ratio (particles/µg protein), which evaluates the overall efficiency of the washing process. As shown in Figure 3(g), the purity ratio significantly improved with washing, increasing by 98% and 160% after one and two washing steps, respectively, compared to the unwashed condition. In conclusion, while the washing step is optional, it is highly recommended when greater purity is required, as it can increase purity by nearly 160% with minimal particle loss.

### 2.4 Scaling up the ExoFilter

To scale up the ExoFilter, we investigated the effect of varying vacuum pressures on the diluted plasma sample 500 mL processing time and particle concentration with the bottle-top type of ExoFilter (Figure 4(a)). Specifically, a 5 mL plasma sample was diluted 1:100 in PBS, yielding a total volume of 500 mL. This diluted sample was then eluted with 100 mL of 1 M NaCl. For the elution step, a volume equivalent to one-fifth of the sample volume was applied as the elution buffer. Pressures ranged from 0 kPa (atmospheric pressure) to 68 kPa. As shown in Figure 4(b), at 0 kPa, processing 500 mL of the sample took 11 min. Increasing the vacuum pressure to 2 kPa reduced the processing time to 2.5 minutes. At 20 kPa, the time further decreased to 80 seconds, and at 68 kPa, the fastest processing time of 40 seconds was achieved for 500 mL (750 mL/min). Regardless of the operating pressure, the EV isolation performance, measured as particle concentration per milliliter, remained consistent across all tested pressures (0, 2, 20, and 68 kPa), as shown in Figure 4(c). This suggests that the ExoFilter system is robust and maintains its efficiency in EV isolation even when scaled up to higher vacuum pressures, ensuring reliable particle recovery without significant variation in concentration.

**Figure 4.**
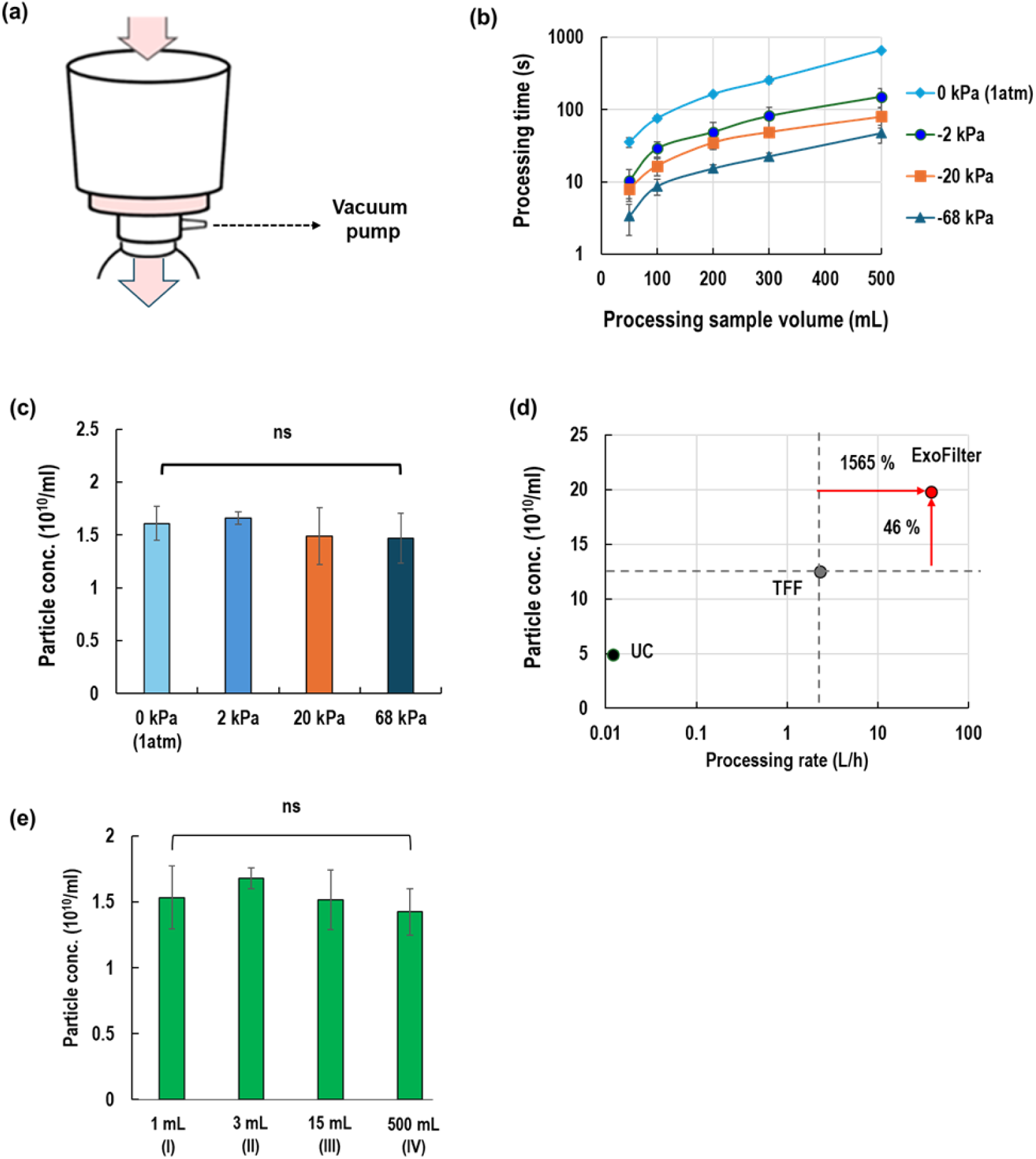
Scaling up the ExoFilter system for large-volume EV isolation from plasma (5 mL) diluted to a final volume of 500 mL (1/100 ratio). (a) Processing times for various sample volumes under different vacuum pressures (0 to 68 kPa), (b) Particle concentrations with varying vacuum pressure, (c) Particle concentration vs. processing rate for UC, TFF and ExoFilter, (d) Comparison of EV isolation yields among three spin-column types and a bottle-top filter type (ExoFilters), (e) Comparison of particle concentrations with varying sample volumes.

Figure 4(d) presents a comparison of particle concentration and processing rate between the ExoFilter and other conventional methods, such as ultracentrifugation (UC) and tangential flow filtration (TFF). The results demonstrate that ExoFilter significantly surpasses both methods. Specifically, the particle concentration obtained with ExoFilter is considerably higher, with a 46% increase over TFF. In this case, ExoFilter can process 500 mL of sample in 40 seconds (750 mL/min or 45 L/h), whereas TFF takes 63 seconds to process just 50 mL (48 mL/min or 2.8 L/h). Therefore, the processing rate of ExoFilter is substantially faster, showing a remarkable 15.5-folds improvement over TFF. This highlights ExoFilter’s superior efficiency in both particle recovery and processing speed, emphasizing its suitability for high-throughput and large-scale EV isolation applications.

The ExoFilter system provides a versatile solution for isolating extracellular vesicles (EVs) from varying sample volumes (1 mL, 3 mL, 15 mL, and 500 mL). The first three are spin column types, and the last one is a bottle-top filter. The operating conditions applied were centrifugation at 5000 × g for 1 minute for the spin-column types and 68 kPa of vacuum pressure for the bottle-top type. We compared the performance of EV isolation using four different types of ExoFilter in Figure 4(e). As long as the appropriate sample volume is applied based on the filter size, nearly identical particle concentrations can be obtained regardless of the filter size and operating mode. These results indicate that the large-scale bottle-top ExoFilter performs comparably to the smaller spin-column types in terms of particle extraction concentration, demonstrating that the system can be effectively scaled up without compromising efficiency.

### 2.5 Performance comparison across different methods using plasma

To evaluate the efficiency of the ExoFilter method, we compared it with other conventional methods, including UC, TFF, ExoQuick, and ExoPAS. Blood plasma was selected as the sample for this study due to its common use in EV extraction, but it presents challenges because of its high protein content. In this experiment, we used the same plasma samples, but the volumes varied according to the requirements of each method: 1 mL for UC, 10 mL for TFF, 1 mL for ExoQuick, 1 mL for ExoPAS, and 200 mL for ExoFilter. To compare each method, we standardized the final concentration ratio to 1:5 during elution.

Figure 5(a) shows the average particle size obtained by each method. The ExoFilter method yielded EVs with an average size of 141 nm, comparable to the sizes obtained using other methods, which ranged from 150 nm to 201 nm. However, the particle concentration achieved by ExoFilter was significantly higher, reaching 18.2 × 10^10^ particles/mL, representing a 46% increase compared to TFF (Figure 5(b)). While ExoQuick and ExoPAS exhibited higher concentrations, these methods likely include protein aggregates and other contaminants that could be mistakenly counted as nano-sized particles, resulting in overestimation.To compensate for the limitations of NTA, we performed RT-PCR analysis to detect specific miRNAs within the EVs. To ensure the amplified miRNA is EV-derived, RNase was added to the extracted EV buffer to remove any potential circulating RNA. Figure 5(c) compares the Ct values of miR-142 and let-7a across the five methods. Surprisingly, the RT-PCR results showed that ExoFilter not only yielded nearly equivalent Ct values to precipitation-based methods like ExoQuick and ExoPAS but also demonstrated significant differences in Ct values of 4.8 and 3.4 compared to TFF. This Ct difference corresponds to approximately a 10-to 20-fold increase in EV concentration.

**Figure 5.**
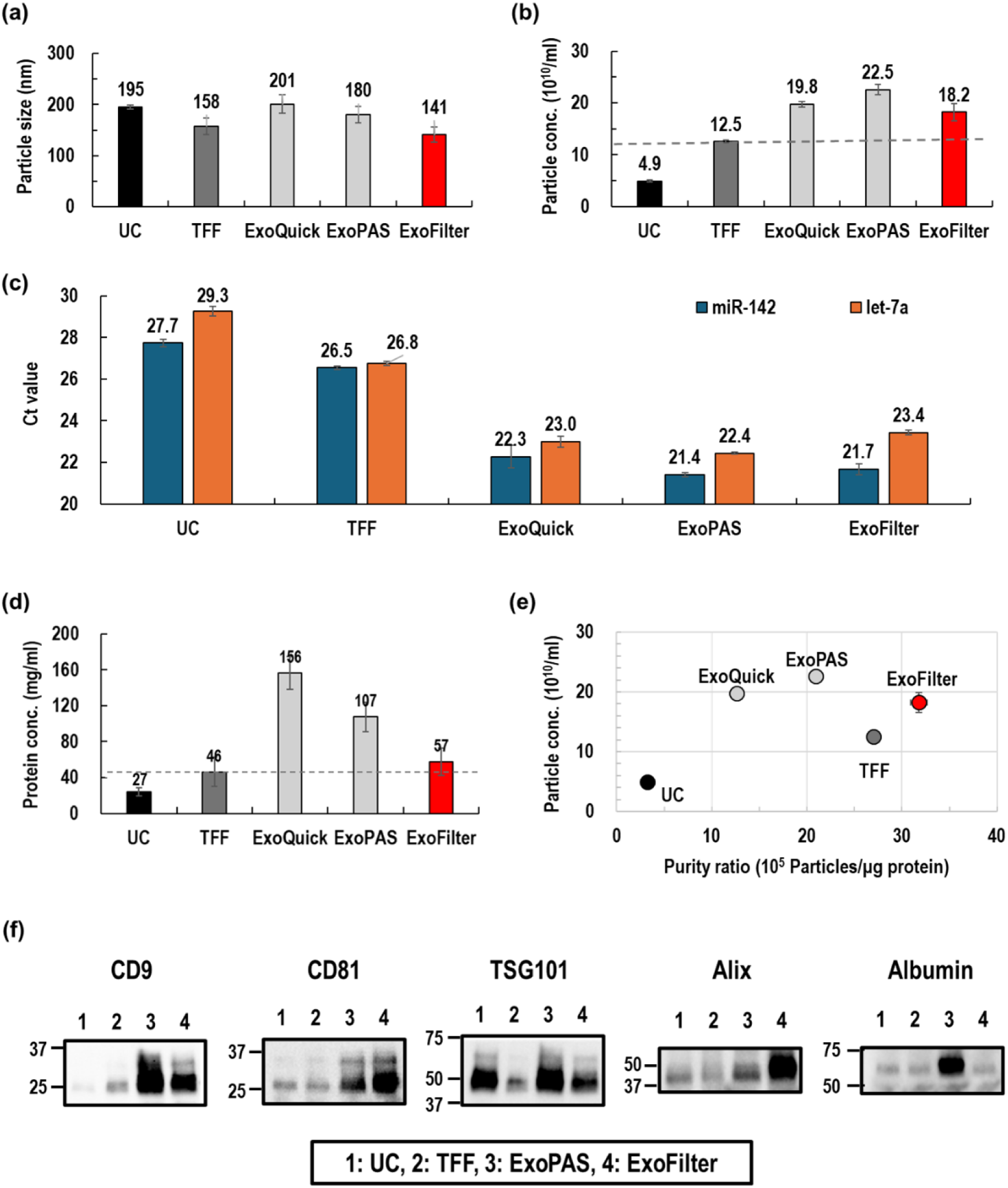
Comparison of EV isolation methods (UC, TFF, ExoQuick, ExoPAS, ExoFilter) using an undiluted plasma sample. (a-b) Mean particle size and particle concentration of isolated EVs, (c) EV-driven miRNAs quantification, (d) Protein concentration of isolated EVs, (e) Purity ratio vs. particle concentration, (f) Western blot for protein markers of EVs and plasma protein marker (albumin).

The protein concentration of the isolated EVs was measured using the BCA assay. Figure 5(d) shows that the ExoFilter method resulted in a protein concentration of 57 µg/mL, which is significantly lower than the concentrations obtained by ExoQuick (156 µg/mL) and ExoPAS (107 µg/mL). This demonstrates the effectiveness of the ExoFilter in reducing protein contamination. By combining the data from particle concentrations and protein concentrations, the purity ratio (particles/µg-protein) was calculated to assess the overall efficiency of EV isolation. Figure 5(e) shows that the ExoFilter demonstrated approximately a 46% increase in particle concentration and a 17% improvement in the purity ratio compared to TFF. Furthermore, ExoFilter achieved the highest purity ratio, significantly outperforming the other methods. This underscores ExoFilter’s superior capability in isolating high-purity EVs from large plasma samples.

In the Western blotting analysis presented in Figure 5(f), the ExoFilter method demonstrated comparable EV protein marker band intensities and thickness to the ExoPAS method. Additionally, ExoFilter exhibited stronger bands than both UC and TFF, indicating higher extraction efficiency of EVs. Notably, the albumin band for the ExoFilter was fainter compared to other methods, suggesting lower contamination and higher purity in EV extraction. The full Western blot image corresponding to Figure 5(f) is available in Figure S1.

### 2.6 Performance comparison for various biofluids

We extended our analysis to determine whether the ExoFilter method could isolate EVs from various biofluids, including CCM, urine, and saliva, in comparison with other conventional methods. For CCM (Figures 6(a-c)), ExoFilter demonstrated a 41% higher particle concentration than TFF and achieved a 30% lower protein concentration, resulting in a 120% higher purity ratio compared to TFF. In fact, ExoFilter exhibited the highest purity ratio of 4.0 × 10^6^ particles per µg protein, significantly outperforming UC, TFF, ExoQuick, and ExoPAS.

**Figure 6.**
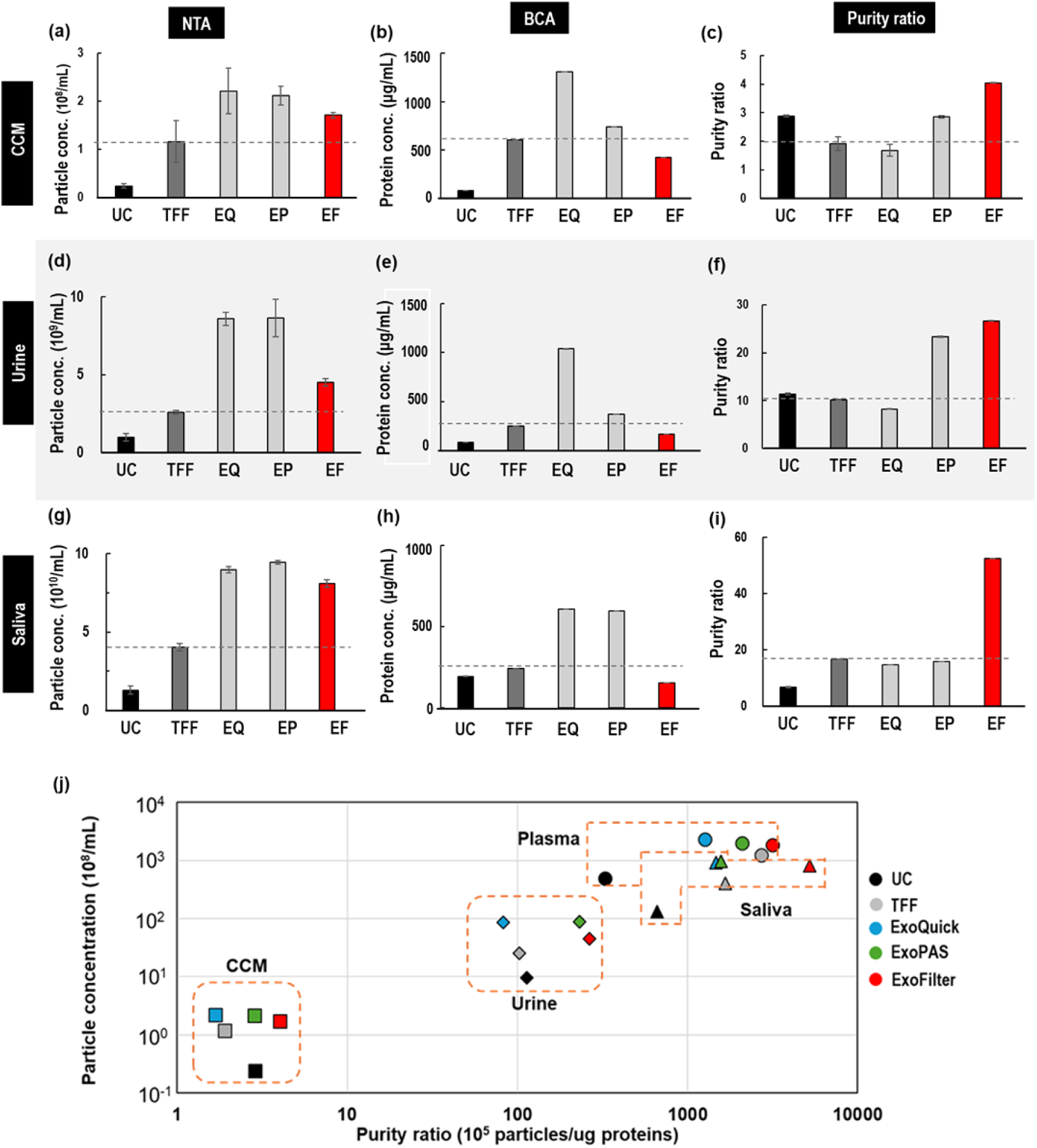
Comparison of EV isolation methods (UC, TFF, ExoQuick, ExoPAS, ExoFilter) across various biofluids. (a-c) Particle concentration, protein concentration and purity ratio of isolated EVs in a CCM sample. (d-f) Particle concentration, protein concentration and normalized purity ratio of isolated EVs in a urine sample. (g-i) Particle concentration, protein concentration and normalized purity ratio of isolated EVs in a saliva sample, (j) Particle concentration versus purity ratio across various biofluids, comparing five different isolation methods.

For urine samples (Figures 6(d-f)), ExoFilter showed superior performance, with a 73% increase in particle concentration compared to TFF and a 162% improvement in purity ratio compared to UC. Additionally, ExoFilter achieved the highest purity ratio among all methods, including UC, TFF, ExoQuick, and ExoPAS. Similarly, in saliva samples (Figures 6(g-i)), ExoFilter maintained a high particle concentration and achieved a significantly lower protein concentration (154 µg/mL), resulting in a 214% higher purity ratio compared to TFF, making it the best-performing method among all those tested.

This study presents an interesting result in Figure 6(j), which illustrates the relationship between particle concentration and purity ratio for various biofluid samples using different extraction methods, including ExoFilter. A key finding from this analysis is that each sample type exhibits distinct characteristics in terms of particle concentration and purity ratio. Specifically, plasma and saliva samples demonstrated the highest particle concentrations and purity ratios across all methods, while urine and CCM samples showed lower values. This variability in extraction efficiency and purity across different sample types underscores the importance of selecting the appropriate extraction method to optimize both yield and purity for each biofluid.

Overall, these results confirm that the ExoFilter method is highly effective for isolating EVs from a variety of biofluids, providing higher particle concentrations and lower protein contamination, leading to superior purity ratios compared to conventional methods. This highlights the versatility and robustness of the ExoFilter system for EV isolation across different biofluid types.

## 3. Discussion and conclusion

Developing a reliable, scalable, and efficient method for isolating high-purity EVs is a key objective in the field, as it would greatly enhance our researches of the EVs in both academic and industrial settings. In this study, we introduce ExoFilter, an innovative platform designed for fast, scalable, and high-performance EV isolation. The ExoFilter method enables the separation of EVs based on electrokinetics while preserving their natural morphology and functionality. Additionally, the scalability of this platform makes it a practical option for large-scale EV isolation from various biofluids. Several studies have reported that when different EV isolation methods are applied sequentially, the unique separation principles of each method result in highly pure EVs at the final stage (Mateescu et al., 2017). The electrokinetic-assisted mesh filtration of ExoFilter can provide a new platform for rapid, large-scale EV isolation.

Our analysis revealed that EVs isolated by ExoFilter exhibited strong signal intensities for specific EV marker proteins and internal nucleic acid markers, while showing weak signals for selected non-EV related proteins. These findings highlight the efficiency and high purity of EV isolation achieved by ExoFilter. Furthermore, ExoFilter can be integrated into existing automated systems (i.e.,, Tanfil-100, Rockers, Taiwan) ensuring a consistent workflow and reproducible results across various sample types, volumes, and EV concentrations. Therefore, ExoFilter offers a convenient and robust approach for the rapid isolation of EVs with superior purity and yield, surpassing traditional EV extraction methods.

In this study, we further explored the electrokinetic-assited filtration mechanism employed by ExoFilter. Figure 7 presents the size and zeta potential distribution of plasma components, including proteins, lipoproteins, and EVs. Although albumin is a small, negatively charged globular protein, Western blot analysis yielded the unexpected result that ExoFilter did not capture albumin (Figure 5f), showing performance nearly equivalent to that of TFF and UC. Albumin depletion in ExoFilter can be explained by the principles of electrokinetic, specifically the interaction between the Coulomb force, which pulls the nanoparticle toward the positively charged substrate, and the velocity field, which drives the nanoparticle along the flow direction.

**Figure 7.**
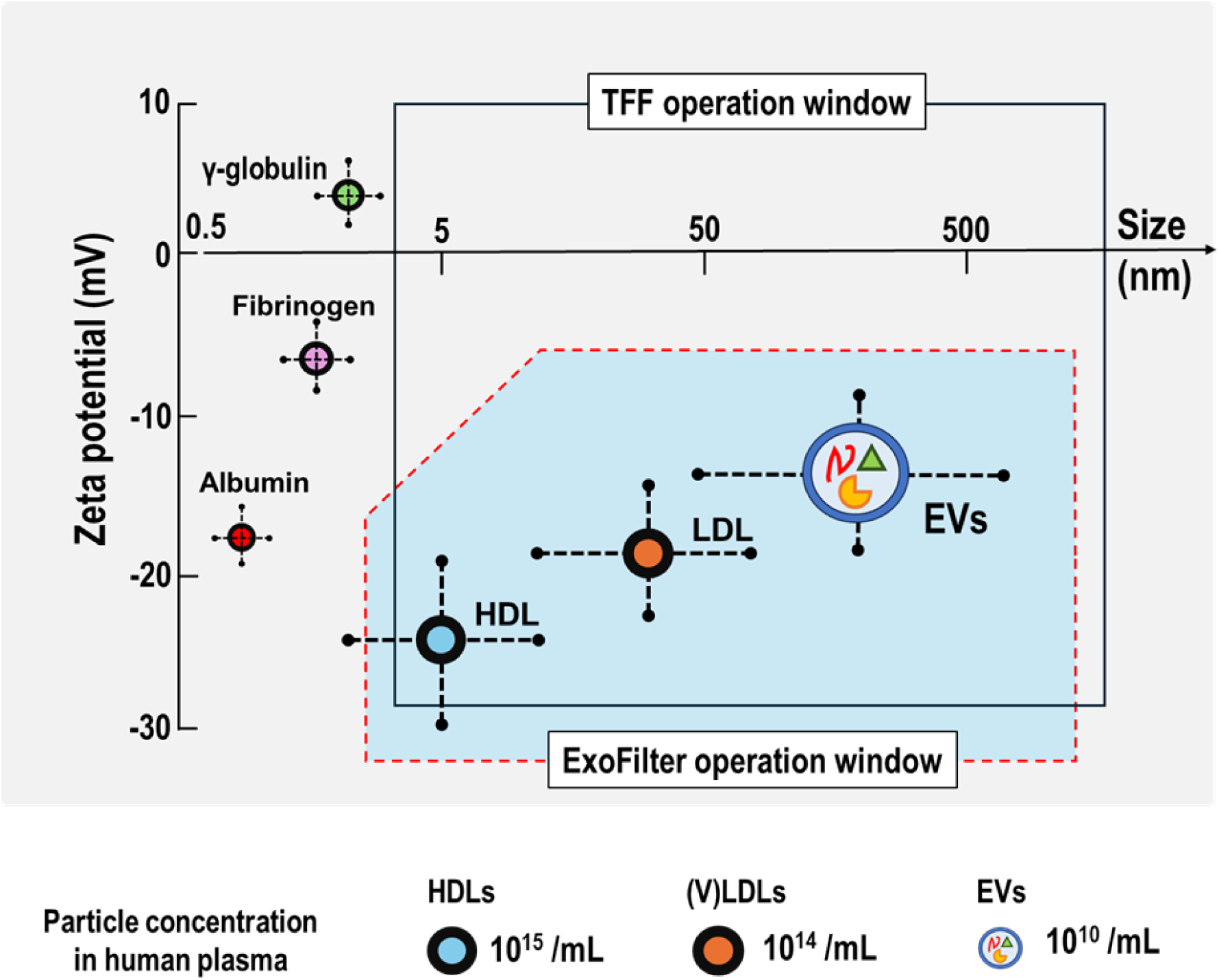
Schematic of the size and zeta potential distribution of plasma proteins, lipoproteins, and EVs. The white box indicates the operation window for TFF based on size cutoff, while the blue box represents that for ExoFilter based on electrokinetic interaction.

The particle’s trajectory is highly influenced by the forces exerted on it within the flow field in Fig. 1(e). First, the horizontal motion of the nanoparticle is governed by the velocity field of the fluid, as shown in Fig. 1(e). Considering the mesh thickness is 100 μm and the constant flow velocity is v = 1 mm/s, the time required for the particle to pass through the mesh is 0.1 seconds. The maximum horizontal displacement (L) during this time would be 0.1 mm due to the short passing time through the mesh. It is important to note that the horizontal displacement of nanoparticles is solely dependent on the fluid velocity under steady flow conditions.

Second, the vertical displacement of a particle is dominated by the Coulomb force resulting from the interaction between its surface charge and the positively charged substrate. This force pulls the particle vertically downward toward the substrate and can be expressed using the equation F_Coulomb_=q⋅σ/(2ɛ_0_) where *q* is a charge of the nanoparticle, *σ* is surface charge density of the substrate, and *ɛ_0_* is permittivity of blood plasma. Albumin size is known as 66.5 kDa or 2.7 nm (Kiselev et al., 2001). Considering size and zeta potential of two nanoparticles, the Coulomb force acting on an EV is approximately 1.29×10^−20^ N, whereas for albumin, it is 3.70×10^−23^ N. The vertical force acting on the EV is nearly 350 times larger than that acting on albumin. If ExoFilter were to operate in a static flow environment, electrokinetic effects would shift to electrostatic interactions, leading to increased albumin capture. This is primarily due to the direct influence of electrical charge characteristics, as previously demonstrated in earlier studies.

In contrast, however, (V)LDL, which has the same level of negative charge as albumin, was not removed through the electrokinetic filtration process of ExoFilter and was captured in similar quantities to EVs (Figure 2(e)). This confirms that (V)LDL cannot be separated by the electrokinetic filtration mechanism in the mesh flow filter, as the size of (V)LDL overlaps significantly with that of EVs. Thus, the operational window for ExoFilter is represented by the blue box in Figure 7, which depicts the pros and cons of the ExoFilter.

One potential avenue for improving EV isolation lies in combining ExoFilter’s charge-based separation method with tangential flow filtration (TFF). As shown in Figure 7, while TFF discards nano-sized particles below a certain threshold by allowing them to pass through the permeate, ExoFilter efficiently isolates EVs based on charge. By integrating these two technologies, it may be possible to achieve higher purity EV isolates, creating a synergistic effect that enhances the overall separation process. This area is under ongoing research, and we are exploring this approach to further improve EV isolation.

Concerns have been raised regarding the potential reduction in the surface functionality of EVs due to the characteristics of the ExoFilter. First, as EVs pass through the nylon mesh multiple times, their surface markers might deteriorate. Additionally, the application of high vacuum pressure (68 kPa) could induce rapid flow rates through micropores, resulting in elevated shear stress that may affect the surface properties of EVs. Second, the charge-based electrostatic attachment and detachment process could potentially impact EV functionality.

The use of 1 M NaCl as an elution buffer introduces hyperosmotic conditions, raising concerns about potential effects on EV characteristics and functionality. As demonstrated in Fig. 2, 1 M NaCl temporarily reduced particle size (104.6 nm) compared to PBS (120.2 nm) and this effect was reversible upon buffer exchange with PBS (122.7 nm). Despite the observed size reduction, cellular uptake and cytotoxicity experiments revealed no adverse effects on EV functionality, even after extended storage at -80°C up to two months. These results indicate that while osmotic pressure may alter EV size, the structural integrity and biological activity of EVs are preserved. Additionally, EVs extracted using ExoFilter from Centella asiatica (CICA) leaves retained their strong wound-healing properties, as shown in Figure S2. This highlights the robustness of EVs under varying buffer conditions and demonstrates the versatility of ExoFilter technology for diverse applications.

The cost-effectiveness and industrial feasibility of ExoFilter present significant advantages in both research and industrial applications. Traditional separation methods, such as size-exclusion chromatography (SEC), rely on expensive materials like microbeads or gels, which often require reuse to remain economically viable. However, reuse is generally restricted in clinical settings, leading to higher costs. In contrast, ExoFilter employs a cost-efficient nylon mesh coated with affordable positively charged substances, significantly lowering production expenses. Its design allows for scalability by adjusting the mesh surface area to accommodate varying sample volumes, while the larger pore size minimizes flow resistance. This enables efficient operation with standard peristaltic or diaphragm pumps, eliminating the need for specialized high-pressure equipment. These features establish ExoFilter as an economical and practical solution for large-scale extracellular vesicle (EV) isolation.

In terms of versatility, ExoFilter excels in both clinical and industrial contexts. The small-volume spin column version is optimized for rapid EV isolation from blood, making it one of the fastest EV extraction techniques available. For industrial-scale applications, ExoFilter offers superior efficiency in processing large sample volumes compared to traditional tangential flow filtration (TFF) systems. Furthermore, recent studies demonstrate that combining ExoFilter with TFF results in synergistic improvements in EV purity and recovery rates. This hybrid approach underscores ExoFilter’s adaptability and scalability, effectively addressing the diverse demands of clinical, research, and industrial environments, and paving the way for broader adoption in EV-related technologies.

## 4 MATERIALS AND METHODS

### 4.1 Preparation of various samples

This study was conducted in accordance with the principles outlined in the Declaration of Helsinki. Plasma samples were obtained from Zen-Bio Inc. (Research Triangle, NC, USA). Cell culture media from umbilical cord mesenchymal stem cells, as well as urine and saliva samples, were collected in sterile containers and subsequently purified through centrifugation. In the EV isolation comparison experiment, the same sample was used for each biofluid. Both types of samples were centrifuged at 3000 × g for 15 minutes to precipitate and remove large particles. The resulting supernatant was then filtered through an 800 nm pore-sized mesh to eliminate particles larger than 800 nm. The processed samples were stored at -80°C until further analysis.

### 4.2 Preparation of cationic polymer coated mesh

Nylon meshes (Lixin Huarun MESH Co., China) were cut into various sizes (d = 7, 11, 22, or 50 mm) using a laser cutter (BEAMO, MIRTECH Korea) and conjugated with cationic materials, such as quaternary ammonium or protamine sulfate. The conjugation process was conducted as follows: The cut nylon meshes (0.5 g) were placed into a Petri dish, and 10 mL of 0.1 M HCl solution was added, followed by a 30-minute incubation. After incubation, the meshes were rinsed three times with deionized (DI) water. Following this, 10 mL of a 2.5% glutaraldehyde solution (Sigma-Aldrich, USA) was added to the petri dish and incubated for 30 minutes. The meshes were then thoroughly rinsed with deionized water to remove any remaining glutaraldehyde. Subsequently, a mixture of 5 mL of 0.1 M EDC solution and 5 mL of 0.1 M NHS solution was added to the dish and incubated for 1 hour. Following the EDC/NHS coupling, the meshes were washed three times with DI water to remove any residual EDC and NHS. The mesh was then conjugated with protamine sulfate salt (Sigma-Aldrich, USA). A solution of 100 mg protamine sulfate in 10 mL of DI water was mixed thoroughly and added to the dish containing the washed mesh. The mixture was incubated for 6 hours at room temperature. After incubation, the protamine sulfate-coated mesh was washed three times with DI water to remove any residual protamine sulfate.

### 4.3 EV isolation

#### 4.3.1 EV isolation procedure using ExoFilter

The charge-based ExoFilter system is capable of processing samples such as plasma, urine, saliva, and cell culture media. It is available in various sizes, designed to handle appropriate volumes based on the characteristics of each sample. The ExoFilter designed for the spin column method can accommodate 1 mL, 3 mL, and 15 mL of samples, while the bottle-top filter version can hold up to 1000 mL. This flexibility allows for tailored experimental design, regardless of the type and volume of the sample, by selecting from a range of available options.

For the bottle-top filter, the sample containing EVs moves to the lower part of the filter by gravity after loading, making it possible to process up to 1000 mL by pouring the sample slowly. To increase the sample flow rate, a vacuum device can be connected to the port on the filter, which further accelerates sample processing (see supplementary video file). In contrast, when using the spin column method, although gravity can assist in moving the sample downward, centrifugation at 5000 × g for 1 minute is optimized to achieve the design objectives. When eluting with 1 M NaCl, it is generally recommended to use an elution volume that is one-fifth of the sample volume. For a 500 mL ExoFilter, 100 mL of elution buffer is sufficient. If further concentration is desired, the elution volume can be reduced, but caution is necessary, taking into account the dead volume of each filter.

#### 4.3.2 EV isolation using ultracentrifuge (UC)

1 mL of various biofluids was mixed with PBS in a 1:3 ratio for EV separation via UC. The mixture underwent sequential centrifugation at 3000 × g for 15 minutes and at 12000 × g for 30 minutes to remove large vesicles such as cell debris. The supernatant was then centrifuged at 120000 × g and 4°C for 2 hours using a high-speed centrifuge (CP100WX; Hitachi, Tokyo, Japan). After the supernatant was discarded, the pellet was resuspended and washed in PBS at 120000 × g and 4°C for 1 hour, then finally resuspended in 200 µL of PBS.

#### 4.3.3 EV isolation by tangential flow filtration (TFF)

For size-based filtration via tangential flow filtration (TFF), we utilized TFF-EVs (Hansa BioMed, Estonia) with an 800 kDa molecular weight cut-off and a fiber pore size of 50 ± 10 nm. Various biofluids, each in a volume of 10 mL, were loaded into the TFF system. The samples were processed by repeatedly passing them back and forth through the TFF device using a syringe until all the liquid had passed through. Subsequently, the washing step was performed by passing 10 mL of PBS back and forth through the system in the same manner. Finally, EV elution was carried out using 2 mL of PBS, which was also passed back and forth through the TFF device to ensure thorough elution of the EVs.

#### 4.3.4 EV isolation with ExoQuick

The ExoQuick exosome precipitation solution (EXOQ5A-1; System Biosciences, Palo Alto, CA, USA) was utilized. A volume of 252 µL of the ExoQuick solution was added to 1 mL of the biofluid sample and incubated for 30 minutes at 4°C. Following incubation, the mixture was centrifuged at 1500 × g for 30 minutes, after which the supernatant was carefully removed, leaving the pellet in the tube. To ensure complete removal of the ExoQuick solution, a second centrifugation at 1500 × g for 5 minutes was performed, and the pellet was then resuspended in 200 µL of PBS.

#### 4.3.5 EV isolation with ExoPAS-2

The ExoPAS-2 method isolates EVs from liquid samples by sequentially utilizing protamine sulfate and PEG (Microgentas, Seoul, Korea). The addition of protamine sulfate causes the negatively charged EVs to form clusters, which are then further aggregated by PEG. Protamine sulfate (1 mg) is first dissolved in deionized water (1 mL) and added to 1 mL of the biofluid sample. PEG 8000 (Sigma-Aldrich, USA) solution (50%, 250 µL) is then added to the mixture, followed by incubation at 4°C for 30 minutes. After incubation, the sample is centrifuged at 3000 × g for 30 minutes, and the supernatant is carefully removed using a pipette. The resulting EV pellet is then resuspended in 200 µL of either PBS or 1 M NaCl solution.

### 4.4 Electron microscopy Images

An anodic aluminum oxide (AAO) membrane mounted in a gasket was utilized for SEM measurements of EVs isolated from plasma. The samples were filtered through the membrane for SEM analysis. Following filtration, the membranes were incubated in a glutaraldehyde solution (Sigma-Aldrich, St. Louis, MO, USA) for 30 minutes. The membranes were then sequentially rinsed with 25%, 50%, 75%, 90%, and 100% ethanol and incubated in a dry oven at 37°C for 2 hours. After coating the membrane with platinum (Pt), the EVs and clusters present on the membranes were observed using an SEM (Quanta 250 FEG; FEI, Hillsboro, OR, USA).

TEM was performed to visualize EVs isolated using various methods, including the ExoFilter. A Caborn Formvar Film-150 copper grid was carefully handled with tweezers, ensuring the sample side was facing upwards, and placed on a Petri dish. Approximately 15 µL of the EV sample was applied to the grid and allowed to adsorb for 1 minute. To avoid contamination, the grid was covered during this incubation period. For negative staining, the grid was positioned vertically at a 90-degree angle, and 1% uranyl acetate was gently applied dropwise with a syringe, allowing the solution to flow downward. Excess staining solution was removed by blotting with filter paper. The grid was then left to air-dry on filter paper. After the drying process, EVs were imaged using a JEM-1400 Flash (JEOL Ltd., Japan) transmission electron microscope. TEM images were captured at an accelerating voltage of 120 kV.

### 4.5 Zeta potential analyzer

The zeta potentials of EVs and several plasma proteins (albumin, γ-globulin, and fibrinogen) were measured using a zeta potential analyzer (Zetasizer Pro; Malvern Panalytical, Malvern, UK). Due to the difficulty in resuspending EVs and plasma proteins in deionized water, a 10-µL sample was diluted with 990 µL of deionized water for the measurements. The cationic nylon mesh was evaluated using a Surpass3 analyzer (Anton Paar GmbH, Austria). For this measurement, a protamine-conjugated mesh with dimensions of 20 mm × 10 mm was utilized.

### 4.6 Nanoparticle tracking analysis (NTA)

Nanoparticle Tracking Analysis (NTA) was conducted using an NS300 instrument and NTA 3.4 Software (NanoSight, Wiltshire, UK). The EV samples were diluted in filtered PBS. For each sample, three 30-second videos were recorded with the camera level set to 14 and the detection threshold set to 11. The video footage of the EVs was captured, and their mean size and concentration were analyzed according to the respective dilution factors.

### 4.7 BCA assay

The Pierce™ BCA Protein Assay Kit (#23225; Thermo Scientific, Waltham, MA, USA) was utilized to assess protein purity. A standard curve ranging from 0 to 2000 μg/mL was established using nine serial dilutions of bovine serum albumin mixed with the working reagent. Each sample and standard point was replicated three times. Samples (100 µL each) were combined with 2.0 mL of the working reagent and incubated at 37°C for 30 minutes. After cooling to room temperature, the absorbance at 562 nm was measured using a spectrometer (DS-11 Fx+; Denovix, Wilmington, DE, USA). The absorbance difference between the sample values and the average absorbance of blank standards was calculated and converted to µg/mL based on the standard curve (Figure S4).

### 4.8 Western blot assay

Proteins for the Western blot assay were extracted from EVs suspended in 200 µL of elution buffer. These proteins were mixed with Laemmli buffer and 2-mercaptoethanol (BioRad, USA) and then heated at 95°C for 10 minutes. Protein samples were separated by gel electrophoresis using SDS-PAGE Mini-PROTEAN® TGX™ Precast Gel (BioRad, USA). For Western blotting, the EV-specific markers CD9, CD81, TSG101, and ALIX were used, along with albumin as a negative marker, in accordance with MISEV 2023 guidelines. The antibodies used for immunoblotting were recombinant anti-CD9 (ab92726, Abcam, Cambridge, UK), recombinant anti-CD81 (ab109201, Abcam, Cambridge, UK), recombinant anti-TSG101 (ab125011, Abcam, Cambridge, UK), recombinant anti-ALIX (ab186429, Abcam, Cambridge, UK), and goat anti-rabbit IgG H&L (ab205718, Abcam, Cambridge, UK). The protein bands were visualized using the ChemiDoc™ XRS+ System (Bio-Rad, CA, USA) with enhanced chemiluminescence (ECL) reagent following antibody incubation.

### 4.9 Immunocapture-based ELISA

Anti-CD9 (R&D Systems, Minneapolis, USA) and anti-Apo B-100 (R&D Systems, Minneapolis, USA), Anti-albumin (R&D Systems, Minneapolis, USA) antibodies were diluted to 5 µg/mL with 10 mM PBS, and 100 µL was dispensed into microwells and fixed at 37°C for 2 hours. The wells were washed with distilled water, and 200 µL of 0.5% casein PBS was added and incubated at 37°C for 1 hour. After another wash, the exosome sample was diluted with 0.5% casein and 0.1% Tween-20 PBS to prepare concentrations of 0, 0.1, 1, 10, and 50 µg/mL, with 100 µL aliquots in each well. The reaction was carried out overnight at 37°C. After washing with distilled water, a biotin-polymerized anti-CD9, Apo-B100, albumin antibody (BioLegend, San Diego, CA) was diluted to 1 µg/mL, and 100 µL was dispensed into microwells for a 1-hour reaction. After washing again, 100 µL of 0.45 µm membrane-filtered streptavidin-poly HRP20 (Fitzgerald, Acton, MA, USA) at 66 ng/mL was added and incubated at 37°C for 1 hour. To generate a signal, 200 µL of HRP substrate solution was dispensed into microwells, followed by a 15-minutes room temperature incubation. To stop the reaction, 50 µL of 2 M sulfuric acid was added to each well, and absorbance at 450 nm was measured using a microplate reader (SPECTROstar Nano, BMG LABTECH, Freiburg, Germany).

### 4.10 miRNA analysis with RT-qPCR

To quantify miRNA markers present in EVs, the extracted RNA was reverse transcribed using the TaqMan MicroRNA RT kit (4366596, Life Technologies, Eugene, OR, USA) in conjunction with TaqMan MicroRNA Assays (4427975, Life Technologies, USA). Specifically, the levels of hsa-let-7a-5p and hsa-miR-142-3p were measured using these assays and TaqMan Universal Master Mix II without UNG (4440040, Life Technologies, Eugene, OR, USA).

### 4.11 Cellular uptake and cytotoxicity test

Human dermal fibroblast (HDF) was a kind gift from Prof. K. M. Park at Incheon National University, Republic of Korea. HDF cells were cultured in Dulbecco’s Modified Eagle’s Medium (DMEM; Corning Inc., USA), supplemented with 10% (v/v) fetal bovine serum (FBS; Gibco, USA) and penicillin-streptomycin (Gibco, USA). All cultures were maintained at 37 °C with 5% CO_2_.

To verify the uptake of EVs by the HDF cell line, EVs were stained with PKH67 green dye (Sigma-Aldrich, Burlington, MA, USA) for 15 minutes at 25°C. The mixture was then filtered through a 100-kDa filter to remove any unbound PKH67 dye. HDF cells were incubated in a medium containing PKH67-labeled EVs at a concentration of 2 × 10^9^ particles/mL. For nuclear staining, Hoechst 33342 dye was added to the culture medium 0.5 mL. The cellular uptake of EVs was observed using a fluorescence microscope (Eclipse Ti2; Nikon, Tokyo, Japan).

To evaluate cell viability, HDF cells were plated in 96-well plates with 0.1 mL of cell culture media per well at a seeding density of 1 × 10⁴ cells/cm² and allowed to adhere for 24 hours. After washing, the cells were exposed to EV-depleted FBS-containing medium and EVs at a concentration of 2 × 10⁹ particles/mL for 72 hours. Cell viability was assessed using a WST-1 assay kit (EZ-Cytox; DoGenBio, Seoul, Korea). The WST-1 reagent was mixed with the culture medium at a 1:10 ratio, and 100 µL of the mixture was added to each well of the 96-well plate. The cells were incubated for 1 hour at 37°C, and absorbance was measured at 450 nm to determine viability.

## 5 Author contributions

KangMin Lee: Formal analysis; investigation; methodology; visualization; writing—original draft. Minju Bae: Formal analysis; investigation; methodology. YongWoo Kim: Formal analysis; investigation; methodology. SoYoung Jeon: Formal analysis; investigation; methodology. Sujin Kang: Formal analysis; investigation; methodology, Wonjong Rhee: Formal analysis; visualization; methodology, Sehyun Shin: Conceptualization; formal analysis; methodology; supervision; visualization; writing—review & editing.

## Supporting information

Demonstration of ExoFilter-BottleTop

## Acknowledgements

This project was conducted with the support of the Alchemist Project of the Korea Evaluation Institute of Industrial Technology (KEIT 20018560/NTIS 1415184668) funded by the Ministry of Trade, Industry & Energy (MOTIE, Korea) and National Research Foundation of Korea (NRF) Grant funded by the Korean Government, MSIP (RS-2023-00207833). The funders had no role in study design, data collection and analysis, decision to publish or preparation of the manuscript.

## 8. Data availability

The main data supporting the results of this study are available within the manuscript and supplementary information files. The raw data files are available for research purposes from the corresponding author upon reasonable request. Source data are provided with this paper.

## 9 Ethics approval and consent to participate

Plasma, urine, and saliva samples were purchased from Zen-Bio Inc. (Research Triangle, NC, USA). These samples were anonymized and did not contain any personal identifiable information, allowing their use without the need for IRB approval.

## Reference

1. Akers, J. C., Gonda, D., Kim, R., Carter, B. S., & Chen, C. C. (2013). Biogenesis of extracellular vesicles (EV): exosomes, microvesicles, retrovirus-like vesicles, and apoptotic bodies. Journal of neuro-oncology, 113, 1–11.

2. Cardoso, R. M., Rodrigues, S. C., Gomes, C. F., Duarte, F. V., Romao, M., Leal, E. C., Freire, P. C., Neves, R., & Simões-Correia, J. (2021). Development of an optimized and scalable method for isolation of umbilical cord blood-derived small extracellular vesicles for future clinical use. Stem cells translational medicine, 10(6), 910–921.

3. Deregibus, M. C., Figliolini, F., D’antico, S., Manzini, P. M., Pasquino, C., De Lena, M., Tetta, C., Brizzi, M. F., & Camussi, G. (2016). Charge-based precipitation of extracellular vesicles. International journal of molecular medicine, 38(5), 1359–1366.

4. Fathali, H., Dunevall, J., Majdi, S., & Cans, A.-S. (2017). Extracellular osmotic stress reduces the vesicle size while keeping a constant neurotransmitter concentration. ACS Chemical Neuroscience, 8(2), 368–375.

5. Gimona, M., Pachler, K., Laner-Plamberger, S., Schallmoser, K., & Rohde, E. (2017). Manufacturing of human extracellular vesicle-based therapeutics for clinical use. International journal of molecular sciences, 18(6), 1190.

6. Kapoor, K. S., Harris, K., Arian, K. A., Ma, L., Zancanela, B. S., Church, K. A., McAndrews, K. M., & Kalluri, R. (2024). High throughput and rapid isolation of extracellular vesicles and exosomes with purity using size exclusion liquid chromatography. Bioactive Materials, 40, 683–695.

7. Kim, H., & Shin, S. (2021). ExoCAS-2: rapid and pure isolation of exosomes by anionic exchange using magnetic beads. Biomedicines, 9(1), 28.

8. Kim, J., Lee, H., Park, K., & Shin, S. (2020). Rapid and efficient isolation of exosomes by clustering and scattering. Journal of clinical medicine, 9(3), 650.

9. Kiselev, M., IuA, G., Dobretsov, G., & Komarova, M. (2001). Size of a human serum albumin molecule in solution. Biofizika, 46(3), 423–427.

10. Lee, M., Choi, W., & Lim, G. (2023). Electrokinetic-assisted filtration for fast and highly efficient removal of microplastics from water. Chemical Engineering Journal, 452, 139152.

11. Momen-Heravi, F., Getting, S. J., & Moschos, S. A. (2018). Extracellular vesicles and their nucleic acids for biomarker discovery. Pharmacology & therapeutics, 192, 170–187.

12. Nawaz, M., Shah, N., Zanetti, B. R., Maugeri, M., Silvestre, R. N., Fatima, F., Neder, L., & Valadi, H. (2018). Extracellular vesicles and matrix remodeling enzymes: the emerging roles in extracellular matrix remodeling, progression of diseases and tissue repair. Cells, 7(10), 167.

13. Paganini, C., Capasso Palmiero, U., Pocsfalvi, G., Touzet, N., Bongiovanni, A., & Arosio, P. (2019). Scalable production and isolation of extracellular vesicles: available sources and lessons from current industrial bioprocesses. Biotechnology Journal, 14(10), 1800528.

14. Petga, M. A. D., Taylor, C., Macpherson, A., Dhadi, S. R., Rollin, T., Roy, J. W., Ghosh, A., Lewis, S. M., & Ouellette, R. J. (2024). A simple scalable extracellular vesicle isolation method using polyethylenimine polymers for use in cellular delivery. Extracellular Vesicle, 3, 100033.

15. Pitt, J. M., Kroemer, G., & Zitvogel, L. (2016). Extracellular vesicles: masters of intercellular communication and potential clinical interventions. The Journal of clinical investigation, 126(4), 1139–1143.

16. Raposo, G., & Stoorvogel, W. (2013). Extracellular vesicles: exosomes, microvesicles, and friends. Journal of Cell Biology, 200(4), 373–383.

17. Stahl, P. D., & Raposo, G. (2018). Exosomes and extracellular vesicles: the path forward. Essays in biochemistry, 62(2), 119–124.

18. Stam, J., Bartel, S., Bischoff, R., & Wolters, J. C. (2021). Isolation of extracellular vesicles with combined enrichment methods. Journal of Chromatography B, 1169, 122604.

19. Théry, C., Witwer, K. W., Aikawa, E., Alcaraz, M. J., Anderson, J. D., Andriantsitohaina, R., Antoniou, A., Arab, T., Archer, F., & Atkin-Smith, G. K. (2018). Minimal information for studies of extracellular vesicles 2018 (MISEV2018): a position statement of the International Society for Extracellular Vesicles and update of the MISEV2014 guidelines. Journal of extracellular vesicles, 7(1), 1535750.

20. Vader, P., Mol, E. A., Pasterkamp, G., & Schiffelers, R. M. (2016). Extracellular vesicles for drug delivery. Advanced drug delivery reviews, 106, 148–156.

21. Welsh, J. A., Goberdhan, D. C., O’Driscoll, L., Buzas, E. I., Blenkiron, C., Bussolati, B., Cai, H., Di Vizio, D., Driedonks, T. A., & Erdbrügger, U. (2024). Minimal information for studies of extracellular vesicles (MISEV2023): From basic to advanced approaches. Journal of extracellular vesicles, 13(2), e12404.

22. Wiklander, O. P., Brennan, M. Á., Lötvall, J., Breakefield, X. O., & El Andaloussi, S. (2019). Advances in therapeutic applications of extracellular vesicles. Science translational medicine, 11(492), eaav8521.

